# Complex behavioral manipulation drives mismatch between host and parasite diversity

**DOI:** 10.1101/001925

**Authors:** Fabricio B. Baccaro, João P. M. de Araújo, Harry C. Evans, Jorge L. P. Souza, William E. Magnusson, David P. Hughes

## Abstract

Parasites and hosts are intimately associated such that changes in the diversity of one partner are thought to lead to changes in the other. We investigated this linked diversity hypothesis in a specialized ant-*Ophiocordyceps* system in three forests across 750 km in Central Amazonia. All species belonging to the fungal genus *Ophiocordyceps* associated with ants have evolved some degree of behavioral control to increase their own transmission, but the leaf-biting behavior is the most complex form of host manipulation. Such a system requires control of the mandibular muscles and a distinct shift in behavior, from climbing vegetation to walking on leaves to rasping leaf veins in the seconds before death. The need to induce complex behavior may limit host availability and represent a constraint on parasite diversity. The consequence for community structure is that complex behavioral manipulation leads to a mismatch between ant hosts and the diversity of their fungal parasites.

## Introduction

Species diversity varies considerably between habitats and regions [1], and the factors driving such heterogeneity typically depend on the scale of the analysis [2]. At the local level, interactions among species are known to play an important role in structuring communities [3]. This is especially the case when the interacting species occupy different trophic levels, which leads to a stronger link between the diversity of consumers and the diversity of resources [4–6]. Parasite-host interactions are examples of such trophic effects. Parasites tend to be host specific with hosts serving as both the habitat and the dispersal agents for parasites [7]. This implies that changes in host abundance often lead to changes in parasite abundance. Such specificity is considered to lead to arms races that promote overall diversity at the community level [8].

Within the framework of linked diversity in host-parasite systems, the sub-set of parasites that manipulate behavior has not been considered. The effect of parasites on their hosts is not only to reduce host fitness but in some cases also involves a manipulation of host behavior that directly increases parasite fitness [9]. In these cases, other constraints acting on the parasite related to its need to control behavior as a life-history strategy may affect the coupling of diversity across scales. The interaction between ants and the ascomycete fungus *Ophiocordyceps* provides a convenient model for understanding the roles of behavioral manipulation on patterns of host and parasite diversity. Ants infected by *Ophiocordyceps* species die in specific locations outside the nest where the microenvironment is ideal for fungal sporulation and subsequent dispersal to new hosts [10–12]. Dying outside the nest is considered adaptive for the fungus, because it avoids the cleaning behavior of ant workers that may prevent the completion of the fungal lifecycle inside the colony [10]. Depending on which fungal species is involved, infected ants may die attached to stems (Fig. 1C), buried in the leaf-litter, attached to tree bark (Fig. 1D) or biting leafs (Fig. 1E). Biting leaf veins or leaf tissue is the most complex form of manipulation and maintains the ant in situ after death giving the fungus the necessary 24-48 hours to grow adhesive mycelia that bind the ant to the plant [11]. This behavioral manipulation is ancient with vein biting occurring at least since the Eocene [12]. Recent evidence has shown that this host-parasite relationship is highly specific with each host species examined having its own specific parasite species [13].

**Figure 1.**
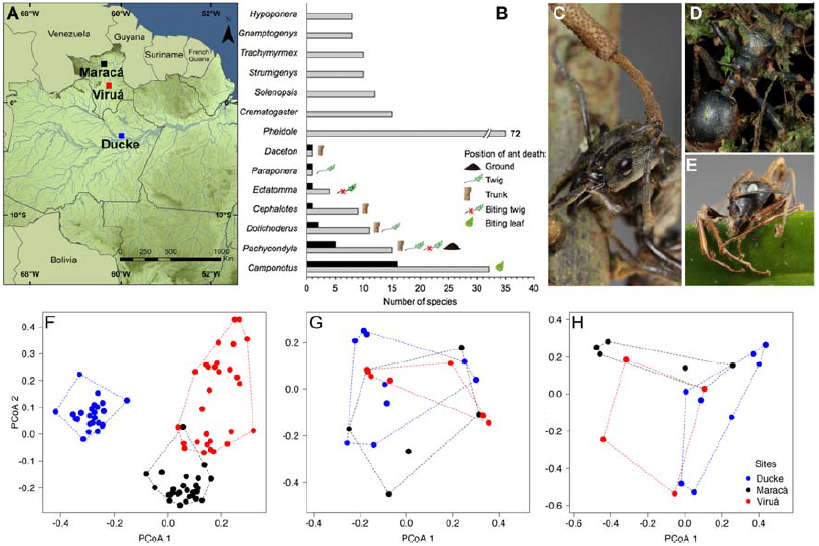
(A) Map of study area. (B) Relative infection levels by ant genera showing where the ants died and the seven more specious non-infected genera sampled in 27 plots (note that *Pheidole* bar is at different scale). Black bar shows the number of infected species and gray bars the number of non-infected species. (C) *Pachycondyla inversa* infected by *Ophiocordyceps kniphofioides* var. *ponerinarum* attached to a stem. (D) *Cephalotes atratus* killed by *O. kniphofioides* var. *kniphofioides* buried in the mosses of a tree trunk. (E) *Camponotus atriceps* parasitized by *O. unilateralis s.l.* biting a leaf edge. PCoA ordination plots indicating (F) the differences in species composition among the three sites using all data, (G) congruence in species composition of all infected ant species found in 25 out of 27 plots and (H) different assemblage composition of non-biting infected ants. Some plots are stacked in the last two figures, because had the same infected ant species composition.

To explore the linked diversity hypothesis between parasite and host we worked with a large dataset of more than 70,000 samples representing 340 ant species with knoweldge on parasite diversity built up from 2,700 samples collected from three Amazonian sites across a 750km transect (Fig. 1A). We specifically compare the composition of infected and non-infected ant species among sites and how the complex behavioral manipulation by the fungus *Ophiocordyceps* can affect the parasite assemblage structure.

## Materials and Methods

We sampled ants and their fungal parasites in three Amazonian forests. Two of them (Maracá Ecological Station, 3^o^ 22’N, 6^o^ 127’W and Viruá National Park, 1^o^ 27’N, 61^o^ 01’W) are situated in forest reserves in Roraima State (extreme North of Brazil). The third (Ducke Reserve, 2^o^ 57’S, 59^o^ 56’ W) is situated 25 km North of Manaus, Central Amazonia (Fig 1A). The sites cover a latitudinal gradient (~ 750 km) in Amazonian forests and encompass wide environmental heterogeneity, including areas of open and dense forests, and areas subject to different degrees of seasonal flooding [14]. We sampled both, parasitized and non-parasitized ants in 9 plots per site covering approximately an area of 9 km^2^. In each plot, we carefully searched for infected ants buried in soil/litter, and attached to vegetation and tree trunks: habitats where the infected ants are most commonly found. The three-dimensional volume sampled per plot was ~ 500 m^3^: 250 m length, 1 m wide and 2 m in height, resulting in 13,500 m^3^ in total. Two persons sampled each plot for at least 1.5 hours (~ 40 hours/person over the three sites); one focusing on all infected ants and the other on non-infected ants belonging to the genera infected by *Ophiocordyceps*. In the Upper Amazon, this and previous research [15] has shown that the following seven ant genera are infected: *Camponotus*, *Cephalotes*, *Daceton*, *Dolichoderus*, *Ectatomma*, *Pachycondyla* and *Paraponera* genera. To contrast the assemblage of infected ants with the whole ant community, we used a comprehensive ant survey of 30 plots per site, with over 70,000 samples collected. This survey included 900 1 m^2^ litter samples (Winkler sacks), 900 pitfall-traps and 900 sardine baits regularly distributed among the three sites to describe the ant assemblage composition (see [14] for additional details).

The data were organized in three matrices: 1) all ant species collected at the three sites, 2) all species from the genera known to be suitable hosts (i.e. *Camponotus*, *Cephalotes*, *Daceton*, *Dolichoderus*, *Ectatomma*, *Pachycondyla* and *Paraponera* genera) and 3) only the species we discovered to be infected. This last category was created because not all species in a genus are infected. To provide further understanding of the role of complex behavioral manipulation on ant community-level patterns, we also constructed matrices of ant species according to the type of manipulation (Fig 1B). We reduced the dimensionality of all matrices using Principal Coordinate Analysis (PCoA) based on the Sørensen dissimilarity index.

Presence/absence data were used to avoid overestimation of species with larger nests. We compared assemblage composition between the three areas using non-parametric multivariate analysis of variance [16]. The statistical significance of each analysis was based on 9,999 Monte Carlo permutations.

## Results

We found that the ant assemblage composition was markedly different between areas (npMANOVA, p < 0.001; Fig 1F). We recorded 343 species from 58 different genera in 11 sub-families. We found very little overlap of species between areas; only 72 out of 343 ant species (~20%) were sampled in each of the three sites. Although we recorded 58 genera of ants only 7 genera contain species that are infected by *Ophiocordyceps* (these are *Camponotus*, *Cephalotes*, *Daceton*, *Dolichoderus*, *Ectatomma*, *Pachycondyla* and *Paraponera*). The assemblage composition of 68 species of ants belonging to these 7 genera also were different between areas (npMANOVA, p < 0.001, Fig S1). However, the assemblage of infected ants did not mirror the community structure; either of all ants from all genera or all ants from the genera that we identified as containing infected species. The assemblage composition of species of ants that are infected by *Ophiocordyceps* was not different between sites (npMANOVA, p = 0.109; Fig 1G). Put another way, despite the fact that ~20% of the ant species were shared among the three locations the infected ant species were similar between these three very geographically and ecologically different sites. We had expected that different areas would have different infected species assemblages reflecting the general pattern of ant diversity across three sites, i.e. linked diversity between hosts and parasites. The linked diversity in the host-parasite system only matched when the infected ants that are manipulated to bite into plant tissue are removed from the analysis (npMANOVA, p < 0.001, Fig. 1H).

## Discussion

To infect an ant worker, *Ophiocordyceps* fungus produces spores that are released onto the forest floor or onto vegetation, over both of which foraging ants move. The spores of this fungus are very large, with a relatively thin cell wall and devoid of pigmentation [i.e. hyaline 17], making them sensitive to dehydration and UV radiation. Therefore, for fungal life cycle completion, an ant must pass, at the correct time of the day, over the spores scattered on the forest floor to be infected. In addition, the parasite must overcome the host population structure to complete their life cycle. Ants are very interactive organisms, and the competition between colonies of the same species at local scale has been frequently demonstrated [18]. Colony-colony competition implies an additional barrier for the transmission between colonies of a specialized parasite. However, the complex behavioral manipulation by *O. unilateralis* seems to circumvent these barriers by creating a relative large minefield areas (~ 30m2, [11]) where the foraging ants can be infected by spores. This is achieved by very high densities of manipulated/killed hosts in a small area known as graveyards (Pontop et al). Virtually, in all the plots where we found species of ants known to be a host to *O. unilateralis* complex we also found ants infected by fungi within the *O. unilateralis* complex. And over the 750 km range the same group of *Camponotus* species were infected despite those areas having limited overlapp in *Camponotus* species asssemblage (Fig 2). Other groups of *Ophiocordyceps* that infect ant species with large colonies, such as the ant genera *Cephalotes* and *Dolichoderus* (both included in this study) also create graveyards, but in smaller areas (personal observations). In the latter case, the dead ants died attached in one tree trunk [17], and sites without any infected ants were more common (Fig 2).

**Figure 2.**
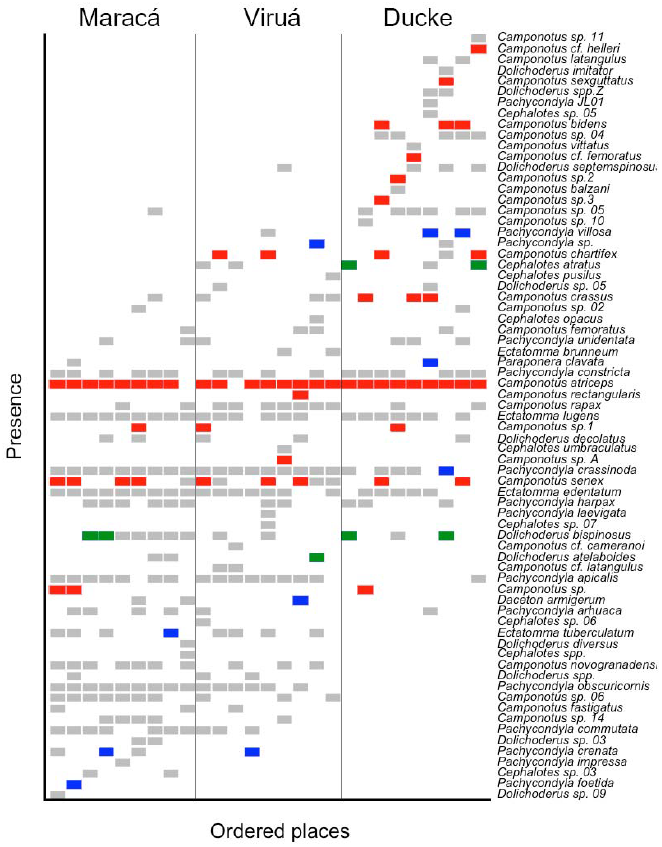
Distribution of ant species ordered by the first axis of PCoA analysis in study plots at Maracá, Viruá and Ducke sites. Occurrence of uninfected ants are in gray. Plots where ant species were infected by *unilateralis* complex (biting plant tissue) are showed in red. Blue and green bars represent plots were ants infected by *australis* and *kniphofioides* complex were found, respectivetly. In the later case, the infected ants were found on litter and buried in tree trunks.

Biting behavior requires a control of the mandibular muscles that involves a reduction in muscle organelle abundance [12]. It also requires a distinct shift in behavior in the seconds before biting as infected ants shift from a wandering behavior to rasping of either the major veins or leaf edges. Other complexes of *Ophiocordyceps* cause ants to die on leaves (*O. lloydii*, 17) but the do not cause ants to bite into the plant tissue. We suggest that the nature of complex manipulation and the necessary additional control of the host’s phenotype that is entailed limit the potential host range of fungi investing in manipulation. Transmission requires this complex control of behavior, which in turn requires multiple effects at the physiological and neuronal level. The consequence for community structure is that even across large geographical areas, complex behavioral manipulation results in a mismatch between host and parasite diversity patterns. However, despite this apparent constraint the evolution of behavioral manipulation seems to be a successful strategy and ant hosts of *O. unilateralis* group were by far the most abundant hosts we discovered.

## Acknowledgements

We thank R. Loreto and C. de Bekker for their help with the field survey; and A. Beattie and H. McCallum for comments on the manuscript. This work was supported by funds from PSU to DPH by CENBAM and PPBio continuous financial support. F.B.B. receive CNPq (140388/2009-5) and CAPES (BEX 8497/11-7) fellowships. Raw data are available at PPBio web site (http://ppbio.inpa.gov.br/knb/style/skins/ppbio/).

